# Inhibition of SARS-CoV-2 infection (previously 2019-nCoV) by a highly potent pan-coronavirus fusion inhibitor targeting its spike protein that harbors a high capacity to mediate membrane fusion

**DOI:** 10.1101/2020.03.09.983247

**Authors:** Shuai Xia, Meiqin Liu, Chao Wang, Wei Xu, Qiaoshuai Lan, Siliang Feng, Feifei Qi, Linlin Bao, Lanying Du, Shuwen Liu, Chuan Qin, Fei Sun, Zhengli Shi, Yun Zhu, Shibo Jiang, Lu Lu

## Abstract

The recent outbreak of coronavirus disease (COVID-19) caused by SARS-CoV-2 infection in Wuhan, China has posed a serious threat to global public health. To develop specific anti-coronavirus therapeutics and prophylactics, the molecular mechanism that underlies viral infection must first be confirmed. Therefore, we herein used a SARS-CoV-2 spike (S) protein-mediated cell-cell fusion assay and found that SARS-CoV-2 showed plasma membrane fusion capacity superior to that of SARS-CoV. We solved the X-ray crystal structure of six-helical bundle (6-HB) core of the HR1 and HR2 domains in SARS-CoV-2 S protein S2 subunit, revealing that several mutated amino acid residues in the HR1 domain may be associated with enhanced interactions with HR2 domain. We previously developed a pan-coronavirus fusion inhibitor, EK1, which targeted HR1 domain and could inhibit infection by divergent human coronaviruses tested, including SARS-CoV and MERS-CoV. We then generated a series of lipopeptides and found that the EK1C4 was the most potent fusion inhibitor against SARS-CoV-2 S protein-mediated membrane fusion and pseudovirus infection with IC_50_s of 1.3 and 15.8 nM, about 241- and 149-fold more potent than that of EK1 peptide, respectively. EK1C4 was also highly effective against membrane fusion and infection of other human coronavirus pseudoviruses tested, including SARS-CoV and MERS-CoV, as well as SARSr-CoVs, potently inhibiting replication of 4 live human coronaviruses, including SARS-CoV-2. Intranasal application of EK1C4 before or after challenge with HCoV-OC43 protected mice from infection, suggesting that EK1C4 could be used for prevention and treatment of infection by currently circulating SARS-CoV-2 and emerging SARSr-CoVs.

## Introduction

In April of 2018, the World Health Organization (WHO) established a priority list of pathogens, including Middle East respiratory syndrome (MERS), severe acute respiratory syndrome (SARS) and Disease X, a disease with an epidemic or pandemic potential caused by an unknown pathogen^1,2^ (Fig.1a).

**Fig. 1.**
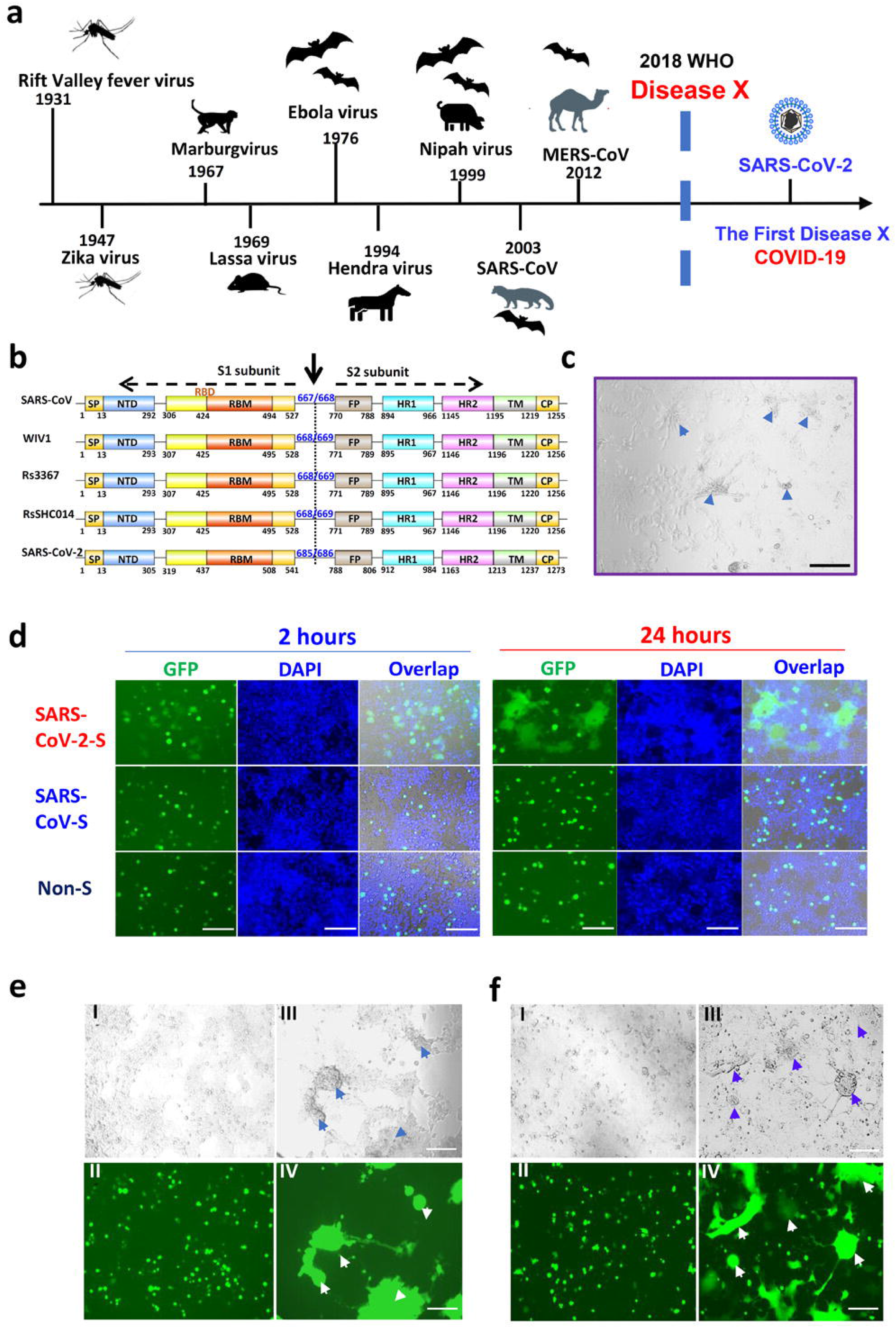
Establishment of SARS-CoV-2 S protein-mediated cell-cell fusion system. **a**. The emerging timeline for highly pathogenic viruses and the proposed Disease X. **b**. Schematic representation of SARS-CoV-2 S protein. Its S1 subunit contains NTD (14-305 aa), RBD (319-541 aa), and RBM (437-508 aa). Its S2 subunit contains FP (788-806 aa), HR1 (912-984 aa), HR2 (1163-1213 aa), TM (1214-1237 aa) and CP (1238-1273 aa). **c**. The formation of syncytium in Huh-7 cells 24 h after SARS-CoV-2 infection, with scale bar of 200 μm. **d**. Images of SARS-CoV- and SARS-CoV-2 S-mediated cell–cell fusion on 293T/ACE2 cells at 2 h (left) and 24 h (right). **e**. SARS-CoV (I-II) and SARS-CoV-2 (III-IV) S-mediated syncytium formation on 293T/ACE2 cells at 48 h. f. SARS-CoV (I-II) and SARS-CoV-2 (III-IV) S-mediated syncytium formation on Huh-7 cells at 48 h. Scale bar equals 400 μm in **d**-**f**.

In late December 2019, an outbreak of pneumonia with an unknown etiology in Wuhan, China was considered as the first Disease X following the announcement by WHO. Shortly thereafter, a novel coronavirus, 2019-nCoV, as denoted by WHO ^3^, was identified as the pathogen causing the coronavirus disease COVID-19 ^4,5^. 2019-nCoV with 79.5% and 96% sequence identity to SARS-CoV and a bat coronavirus, SL-CoV-RaTG13, respectively ^6^, was renamed SARS-CoV-2 by the Coronaviridae Study Group (CSG) of the International Committee on Taxonomy of Viruses (ICTV) ^7^, while, in the interim, it was renamed HCoV-19, as a common virus name, by a group of virologists in China ^8–10^.

As of 24 February 2020, a total of 79,331 confirmed cases of COVID-19, including 2,618 deaths, were reported in China and 27 other countries ^11^, posing a serious threat to global public health and thus calling for the prompt development of specific anti-coronavirus therapeutics and prophylactics for treatment and prevention of COVID-19.

Coronaviruses (CoVs), the largest RNA viruses identified so far, belonging to the *Coronaviridae* family, are divided into 4 genera, α-, β-, δ- and γ-coronaviruses, while the β-coronaviruses are further divided into A, B, C, and D lineages. The seven CoVs that can infect humans (HCoVs) include HCoV-229E and HCoV-NL63 in the α-coronaviruses, HCoV-OC43 and HCoV-HKU1 in the β-coronaviruses lineage A, SARS-CoV and SARS-CoV-2 in the β-coronaviruses lineage B (β-B coronaviruses), and MERS-CoV in the β-coronaviruses lineage C ^6^. To develop specific SARS-CoV-2 fusion inhibitors, it is essential to study the fusion capacity of SARS-CoV-2 compared to that of SARS-CoV. Particularly, the spike (S) protein S2 subunit of SARS-CoV-2, which mediates membrane fusion, has 89.8% sequence identity and 96.9% sequence similarity to those of SARS-CoV, and both of them utilize human angiotensin-converting enzyme 2 (hACE2) as the receptor to infect human cells ^6^. Most importantly, the ACE2-binding affinity of the receptor-binding domain (RBD) in S1 subunit of S protein of SARS-CoV-2 is 10- to 20-fold higher than that of SARS-CoV ^12^, which may contribute to the higher infectivity and transmissibility of SARS-CoV-2 compared to SARS-CoV. However, it is unclear whether SARS-CoV-2 can mediate membrane fusion in a manner that exceeds the capacity of SARS-CoV.

After binding of RBD in S1 subunit of S protein on the virion to the ACE2 receptor on the target cell, the heptad repeat 1 (HR1) and 2 (HR2) domains in its S2 subunit of S protein interact with each other to form a six-helix bundle (6-HB) fusion core, bringing viral and cellular membranes into close proximity for fusion and infection ^13^. Therefore, the 6-HB fusion core structure of SARS-CoV-2 and SARS-CoV S proteins should also be compared in order to investigate the structural basis for membrane fusion mediated by their S proteins and thus set the stage for the rational design of coronavirus fusion inhibitors.

In our previous studies, we designed a pan-coronavirus fusion inhibitor, EK1, targeting the HR1 domains of HCoV S proteins, which proved to be effective in inhibiting infection of 5 HCoVs, including SARS-CoV and MERS-CoV, and 3 SARS-related CoVs (SARSr-CoVs). By intranasal application of this peptide, either pre- or post-challenge with a coronavirus, the treated mice were protected from HCoV-OC43 or MERS-CoV infection, suggesting that this peptide has prophylactic and therapeutic potential against SARS-CoV-2 infection^14^. Indeed, our recent studies have shown that EK1 peptide is effective against SARS-CoV-2 S protein-mediated membrane fusion and PsV infection in a dose-dependent manner ^15^.

In this study, we have shown that SARS-CoV-2 exhibits much higher capacity of membrane fusion than SARS-CoV, suggesting that the fusion machinery of SARS-CoV-2 is an important target for development of coronavirus fusion inhibitors. We have solved the X-ray crystal structure of SARS-CoV-2’s 6-HB core and identified several mutated amino acid residues in HR1 domain responsible for its enhanced interactions with HR2 domain. By conjugating the cholesterol molecule to the EK1 peptide, we found that one of the lipopeptides, EK1C4, exhibited highly potent inhibitory activity against SARS-CoV-2 S-mediated membrane fusion and PsV infection, about 240- and 150-fold more potent than EK1 peptide, respectively. EK1C4 is also highly effective against *in vitro* and *in vivo* infection of some live HCoVs, such as SARS-CoV-2, HCoV-OC43 and MERS-CoV, suggesting potential for further development as pan-CoV fusion inhibitor-based therapeutics and prophylactics for treatment and prevention of infection by the currently circulating SARS-CoV-2 and MERS-CoV, as well as future reemerging SARS-CoV and emerging SARSr-CoVs.

## Results

### The capacity of SARS-CoV-2 S protein-mediated membrane fusion

From the GISAID Platform (https://platform.gisaid.org), we obtained the full-length amino-acid sequence of SARS-CoV-2 (BetaCoV 2019-2020) S protein (GenBank: QHD43416). Through alignment with SARS-CoV and SL-CoVs S proteins, we located the functional domains in SARS-CoV-2 S protein, which contains S1 subunit and S2 subunit with the cleavage site at R685/S686 ^15^. S1 subunit is located within the N-terminal 14–685 amino acids of S protein, containing N-terminal domain (NTD), receptor binding domain (RBD), and receptor binding motif (RBM). S2 subunit contains fusion peptide (FP), heptad repeat 1 (HR1), heptad repeat 2 (HR2), transmembrane domain (TM) and cytoplasmic domain (CP) (Fig. 1b).

Recent biophysical and structural evidence showed that SARS-CoV-2 S protein binds hACE2 with 10-fold to 20-fold higher affinity than SARS-CoV S protein, suggesting the higher infectivity of the new virus ^12^. Unlike other β-B coronaviruses, S protein of SARS-CoV-2 harbors a special S1/S2 furin-recognizable site, indicating that its S protein might possess some unique infectious properties. Indeed, in live SARS-CoV-2 infection, we found a typical syncytium phenomenon naturally formed by infected cells, which is rarely reported in SARS-CoV infection (Fig. 1c). To further explore the special characteristic of SARS-CoV-2 infection, we cloned the S gene into PAAV-IRES-GFP vector and established the S-mediated cell-cell fusion system, using 293T cells that express SARS-CoV-2 S protein and EGFP (293T/SARS-CoV-2/EGFP) as the effector cells, and ACE2/293T cells expressing human ACE2 receptor as the target cells (Fig. 1d and Fig. S1a). After effector cells and target cells were cocultured at 37 °C for 2 h, the fused cells showed at least 2-fold larger size than normal cells and multiple nuclei, and these cells were observed in the SARS-CoV-2 group, but not the SARS-CoV group. After coincubation for 24 h, hundreds of target cells fused together as one big syncytium, containing multiple nuclei (Fig. 1d). Another 24h later, the syncytium grew bigger and could be easily observed under both light and fluorescence microscopy (Fig. 1e). Similar results were observed in the fusion between 293T/SARS-CoV-2/EGFP cells and Huh-7 cells, which naturally express human ACE2 receptor on the cell surface. Their syncytium was obviously formed after coincubation for 48 h, similar to the syncytium formed by live SARS-CoV-2-infected Huh-7 cells (Fig. 1c and 1f). On the contrary, SARS-CoV S protein lacked the ability to mediate the cell-cell fusion under the same conditions (Fig. 1d) based on the required presence of exogenous trypsin to complete membrane fusion in our previous studies. Therefore, compared to SARS-CoV, SARS-CoV-2 S protein showed much more efficiency in mediating viral surface-fusion and entry into target cells ^14^. Meanwhile, no fusion was observed for 293T/EGFP cells without S-expression or 293T cells without ACE2-expression (Fig. 1d and Fig. S1b), confirming that S-receptor engagement is necessary for the S-mediated viral fusion and entry.

### X-ray crystallographic analysis of the 6-HB fusion core formed by HR1 and HR2 domains in S2 subunit of SARS-CoV-2 S protein

Previously, we identified that the 6-HB formed by HR1 and HR2 domains of the S2 subunit plays a very important role in the membrane fusion process mediated by MERS-CoV or SARS-CoV S protein ^16,17^. Similarly, our recent study suggested that HR1 and HR2 in subunit S2 of SARS-CoV-2 also interacted to form coiled-coil complex to support membrane fusion and viral infection^15^ (Fig. 2a and Fig. S2). However, the specific binding characteristics of SARS-CoV-2 6-HB remained to be explored.

**Fig. 2.**
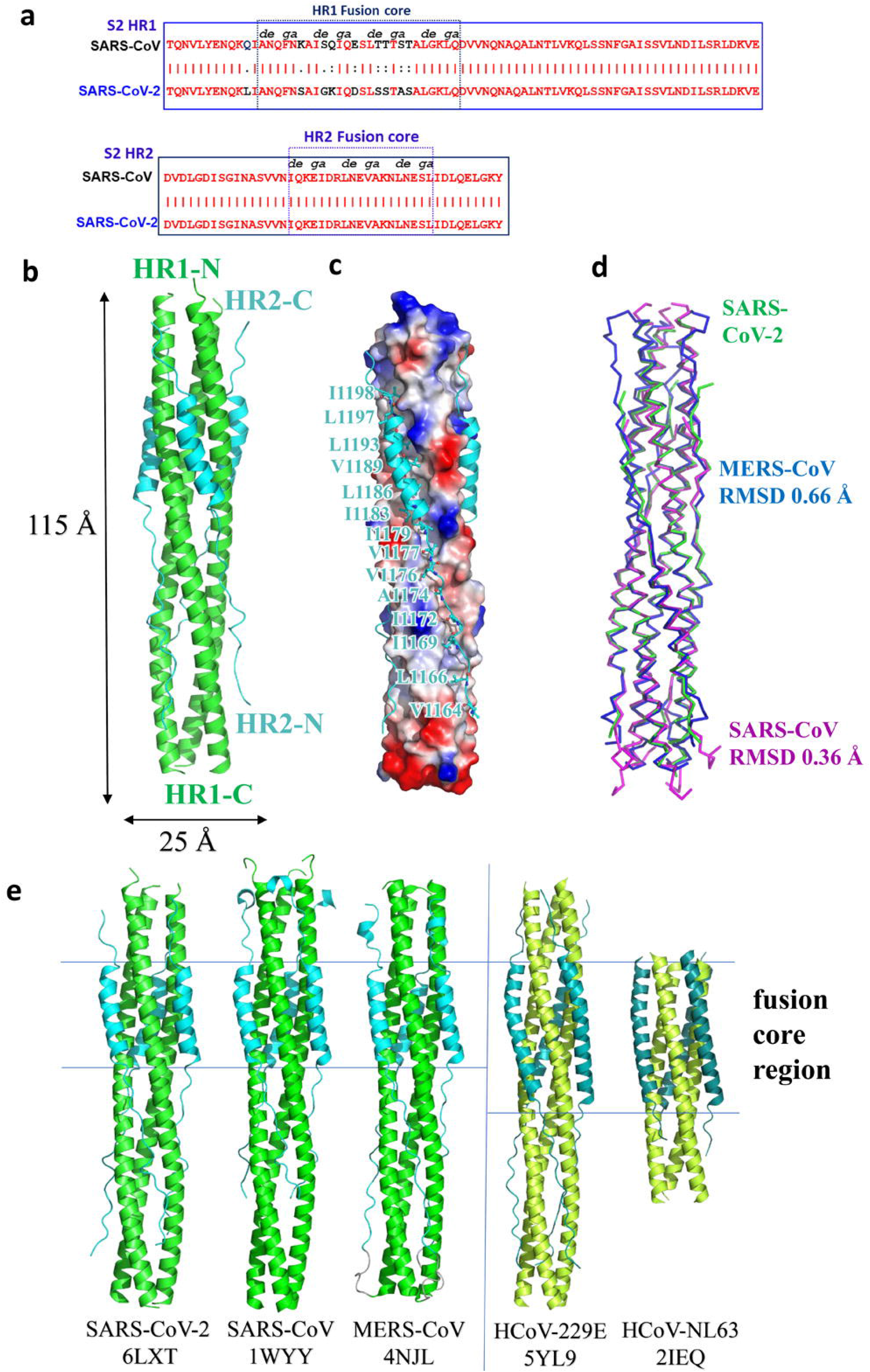
Overall structure of post-fusion 6-HB in SARS-CoV-2. **a.** Sequence alignment of HR1 and HR2 domains in SARS-CoV and SARS-CoV-2. **b**. Structure of SARS-CoV-2 6-HB is shown in cartoon representation with HR1 colored in green and HR2 in cyan. The structural dimensions are indicated in angstroms. **c**. HR1 trimer of SARS-CoV-2 6-HB is shown in electrostatic surface, and HR2 domain is shown in cartoon representation, the important binding residues of which are shown in sticks and labeled. **d**. The superposition of 6-HB structure of SARS-CoV (PDB entry 1WYY), MERS-CoV (PDB entry 4NJL) and SARS-CoV-2 is shown in ribbon. The RMSD between structures is indicated. **e**. The sequence comparison of 6-HB structure of different HCoVs is shown in cartoon representation with different colors for HR1 and HR2. The helical fusion core regions are indicated.

To understand the structural basis of the interactions between HR1 and HR2 regions of SARS-CoV-2, a fusion protein containing the major parts of HR1 (residues 910 to 988) and HR2 (residues 1162 to 1206) with a flexible linker (L6, SGGRGG) in between was constructed for crystallographic study. The crystal structure of HR1-L6-HR2 shows a canonical 6-HB structure with a rod-like shape 115 Å in length and 25 Å in diameter (Fig. 2b). The three HR1 domains form a parallel trimeric coiled-coil center, around which three HR2 domains are entwined in an antiparallel manner. The interaction between these two domains is predominantly a hydrophobic force. Each pair of two adjacent HR1 helices forms a deep hydrophobic groove, providing the binding site for hydrophobic residues of the HR2 domain, including V1164, L1166, I1169, I1172, A1174, V1176, V1177, I1179, I1183, L1186, V1189, L1193, L1197 and I1198 (Fig. 2c). The hydrophobic interactions between HR1 and HR2 are mainly located in the helical fusion core region, which will be discussed later.

The overall 6-HB structure of SARS-CoV-2 is similar to that of other HCoVs with root-mean-square deviation (RMSD) of 0.36 Å to SARS-CoV 6-HB and 0.66 Å to MERS-CoV 6-HB for all the Cα atoms (Fig. 2d). This finding suggested that the overall 6-HB conformation is an important and highly conserved component for these dangerous coronaviruses. When comparing with the 6-HB of other common coronaviruses causing mild respiratory disease, such as 229E and NL63, the SARS-CoV-2 6-HB has a similar overall structure, except for the different length of HR2 helix in the 6-HB. The HR2 domain of 229E or NL63 forms a longer and bending helix to interact with trimeric HR1 core (Fig. 2e). The relationship between the structural difference and the pathogenicity of these HCoVs remains to be elucidated.

According to sequence alignment, the S2 subunits of SARS-CoV-2 and SARS-CoV are highly conserved, with 92.6% and 100% overall homology in HR1 and HR2 domains, respectively. Inside the fusion core region of HR1 domain, there are 8 different residues (Fig. 3a), which may contribute the enhanced interactions between HR1 and HR2 and stabilize 6-HB conformation of SARS-CoV-2 as revealed by crystallographic analysis, compared with those of SARS-CoV. This significant difference has not been observed in other SARS-like viruses, such as WIV1, Rs3367, and RsSHC014. As shown in Figure 3b, the K911 in SARS-CoV HR1 could bind to E1176 in HR2 through a salt bridge 2.9 Å in distance. However, with the Lys-Ser replacement, S929 in SARS-CoV-2 binds to S1196 through a strong hydrogen bond 2.4 Å in distance. In SARS-CoV, Q915 in the HR1 domain does not bind to the HR2 domain. However, with Q-K replacement in the new virus, K933 in the HR1 domain binds to carbonyl oxygen of N1172 in HR2 through a salt bridge 2.7 Å in distance (Fig. 3b). In SARS-CoV, E918 in the HR1 domain binds to R1166 in the HR2 domain through a weak salt bridge 3.7 Å in distance. In SARS-CoV-2, E918 is mutated to D936 and binds to R1185 in the HR2 domain through a salt bridge 2.7 Å in distance (Fig. 3c). In SARS-CoV, K929 in HR1 binds to E1163 in HR2 through a salt bridge 3.2 Å in distance, while T925 is not involved in the interaction. However, when T925 was mutated to S943, it could bind to E1182 in the HR2 domain with a hydrogen bond 2.6 Å in distance, and K947 could also bind to E1182 through a salt bridge 3.0 Å in distance (Fig. 3d). These results suggested that the multiple replacements in the HR1 domain of emerging SARS-CoV-2 virus could enhance the interactions between HR1 and HR2 domain to further stabilize the 6-HB structure, which may lead to increased infectivity of the virus.

**Fig. 3.**
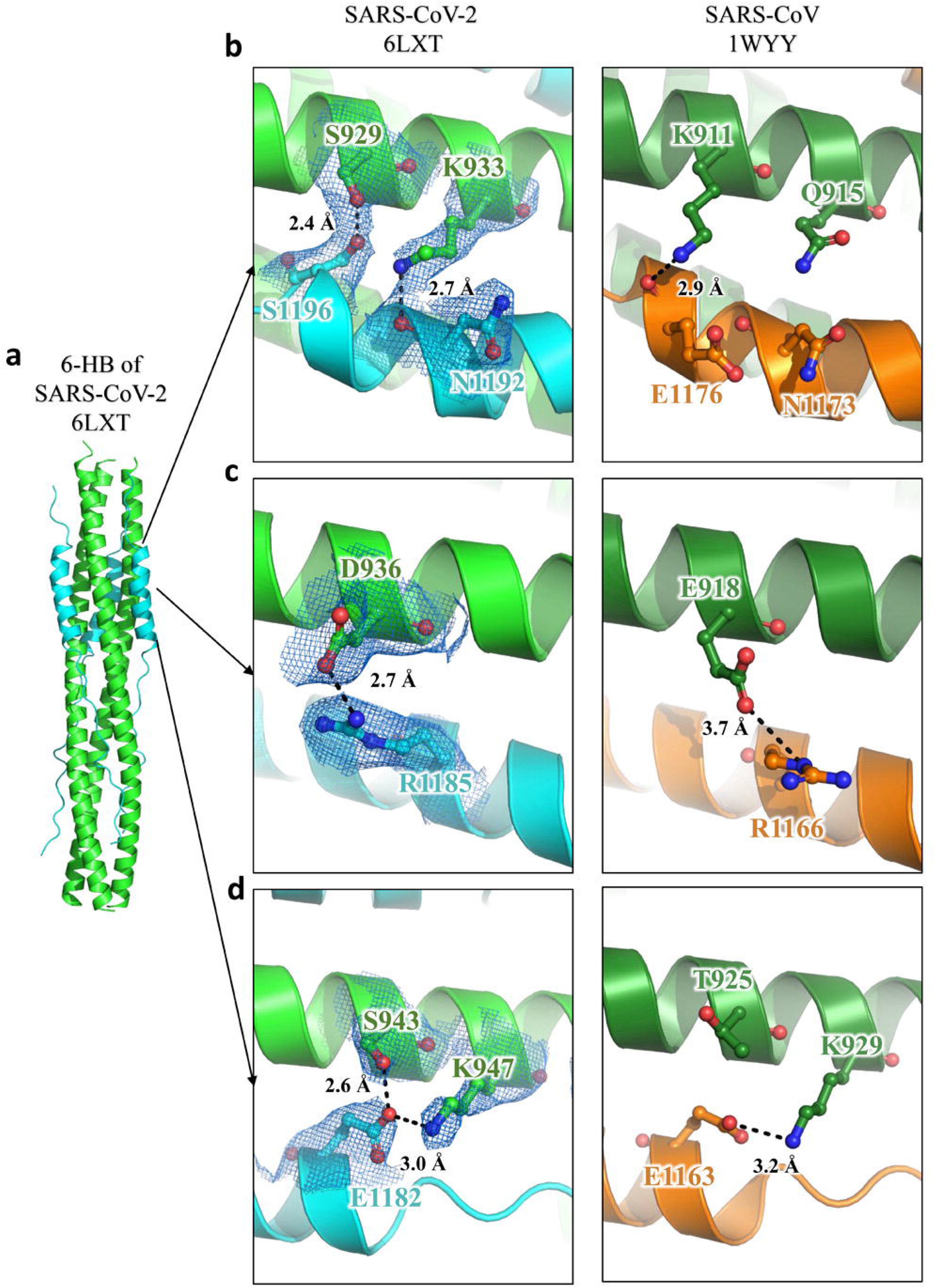
Interaction between HR1 and HR2 of SARS-CoV-2 and SARS-CoV. **a**-**d**. The 6-HB structure of SARS-CoV-2 and SARS-CoV is shown in cartoon representation. The HR1 domain is shown in green for SARS-CoV-2 and forest for SARS-CoV, while the HR2 domain is shown in cyan for SARS-CoV-2 and orange for SARS-CoV. Important residues are shown in sticks and labeled.

### Design and structure-activity relationship (SAR) analysis of lipopeptides with remarkably improved fusion inhibitory activity

Previously, we found that peptide EK1 could disturb viral 6-HB formation and effectively inhibit SARS-CoV-2 PsV infection. However, the potent stability of SARS-CoV-2 6-HB structure might reduce the antiviral efficacy of EK1. Recently, numerous reports have shown that the lipidation strategy can effectively improve the antiviral activity of fusion inhibitory peptides, such as the ant-HIV-1 peptide LP-19 ^18^, and the anti-Nipah virus lipopeptides ^19^. In order to improve the inhibitory activity of EK1, cholesterol (Chol) and palmitic acid (Palm) were covalently attached to the C-terminus of EK1 sequence under the help of a flexible polyethylene glycol (PEG) spacer, and the corresponding lipopeptides EK1C and EK1P were constructed, respectively (Fig. 4a). Both of them could completely inhibit SARS-CoV-2 mediated cell-cell fusion at the concentration of 2.5 μM (Fig. 4b). The inhibitory activity with mean 50% inhibitory concentration (IC_50_) values is 48.1 nM for EK1C and 69.2 nM for EK1P, respectively (Fig. 4c). Meanwhile, the EK1-scrambled peptide showed no inhibitory activity with the concentration up to 5 μM (Fig. 4c). These results strongly suggest that lipidation of EK1 is a promising strategy to improve its fusion-inhibitory activity against SARS-CoV-2 infection, especially, cholesterol-modification.

**Fig. 4.**
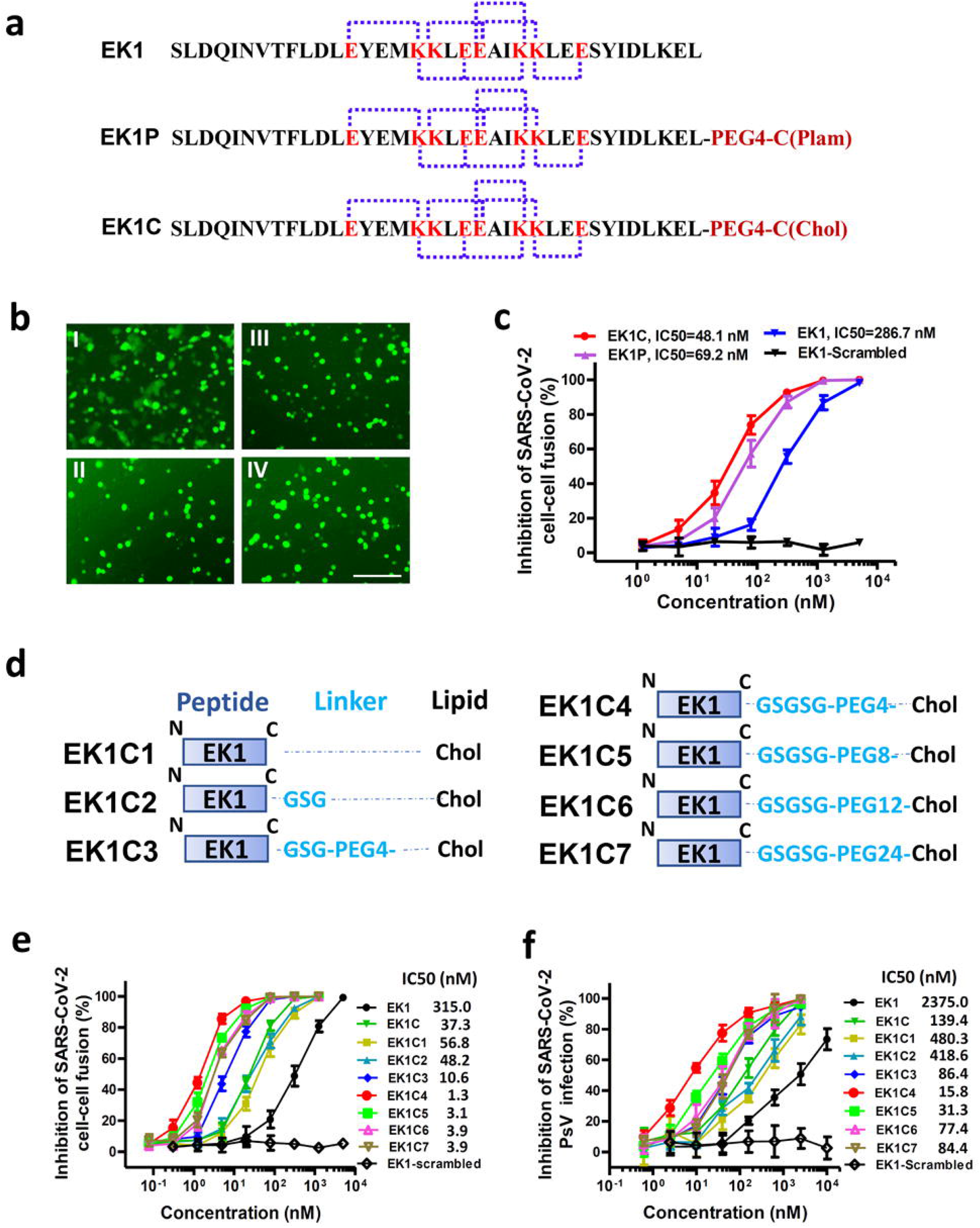
EK1-Lipopeptides showed potent inhibitory activity against SARS-CoV-2 infection. **a**. Amino acid sequences of the designed peptides EK1 and EK1C. The dotted lines represent E–K salt-bridge with i to i + 3, or i + 4 arrangement. **b**. SARS-CoV-2 S protein-mediated cell-cell fusion in the presence of EK1-scramble (I), EK1 (II), EK1C (III), and EK1P (IV) at 2.5 μM (scale bar: 400 μm). **c**. Inhibitory activity of EK1-scramble, EK1, EK1C and EK1P against SARS-CoV-2 S-mediated cell-cell fusion. **d**. Design diagram of EK1-lipopeptides with cholesterol modification, including EK1C1-EK1C7. **e**. Inhibitory activity of EK1-lipopeptides on SARS-CoV-2 S-mediated cell-cell fusion. **f**. Inhibitory activity of EK1-lipopeptides on SARS-CoV-2 PsV infection. Experiments were repeated twice, and the data are expressed as means ± SD (error bar).

On the basis of the structure of EK1C, series of cholesteryl EK1 with multiple linkers were constructed, where the glycine/serine-based linker, *i.e.*, GSG, or PEG-based spacer was employed between EK1 and the cholesterol moiety (Fig. 4d). Compared with EK1C1, EK1C2 and EK1C showed similar inhibitory activities. Strikingly, EK1C3 peptide with both the 3-amino acid linker “GSG” and the PEG4-based spacer, exhibited 4-fold more potency than EK1C1. It is noteworthy that changing “GSG” in EK1C3 to a longer 5-amino acid linker “GSGSG” significantly increased the inhibitory potency of the hybrid molecule, and EK1C4 had IC_50_ value of 1.3 nM, which was 43-fold more potent than EK1C1. These findings indicate that the linker length has a significant effect on the overall activity of lipopeptides.

Comparison of increasing PEG-based arm lengths in EK1C4 shows that inhibitors potency slightly decreased in the cell-cell fusion assay (Fig. 4e). The data suggest that “GSGSG-PEG4” linker was optimal to bridge both parts of the conjugates. Similarly, EK1C4 showed the most potent inhibitory activity against SARS-CoV-2 PsV infection, with IC_50_ value of 15.8 nM, providing 149-fold stronger anti-SARS-CoV-2 activity than that of EK1 (IC_50_=2,375 nM) (Fig. 4f).

### The lipopeptide EK1C4 exhibits the most potent inhibitory activity against membrane fusion mediated by S proteins and entry of pseudotyped coronaviruses

We have previously demonstrated that EK1 could effectively inhibit divergent HCoV infection by targeting the HR1 domains, including α-HCoV and β-HCoV. Here, we further systematically evaluated the broad-spectrum surface-fusion inhibitory activity of EK1C4 on cell-cell fusion mediated by S proteins of divergent coronaviruses, including SARS-CoV, MERS-CoV, HCoV-OC43, HCoV-NL63 and HCoV-229E. Among them, SARS-CoV has the closest relatives to SARS-CoV-2, and its S protein-mediated cell-cell fusion could be effectively inhibited by EK1C4 with IC_50_ of 4.3 nM, which is about 94-fold more active than that of EK1 (IC_50_ = 409.3 nM) (Fig. 5a). Similarly, EK1C4 showed extremely potent fusion-inhibitory activity on MERS-S- and OC43-S-mediated cell-cell fusion with IC_50_ of 2.5 nM and 7.7 nM, which were 95- and 101-fold more potent when compared to EK1, respectively, indicating that EK1C4 could potently and broadly inhibit S protein-mediated cell-cell fusion of various β-HCoVs (Fig. 5b-c). For α-HCoVs, EK1C4 also effectively blocked the fusion process mediated by the S protein of HCoV-229E and HCoV-NL63 with IC_50_ of 5.2 nM and 21.4 nM, respectively, while EK1 showed inhibitory activity of IC_50_ ranging from 207.4 to 751.0 nM (Fig. 5d-e). Moreover, with their potential for human infection, SL-CoVs, including WIV1, Rs3367 and RsSHC014, the fusion process of which is mediated by S protein, could also be significantly prevented by EK1C4 with IC_50_ ranging from 4.3 to 8.1 nM, as well as EK with IC_50_ ranging from 237.0 to 279.6 nM (Fig. 5f-h). As control, the EK1-scrambled peptide showed no inhibitory activity with concentration up to 5 μM in all those coronavirus cell-cell fusion assays (Fig. 5a-h).

**Fig. 5.**
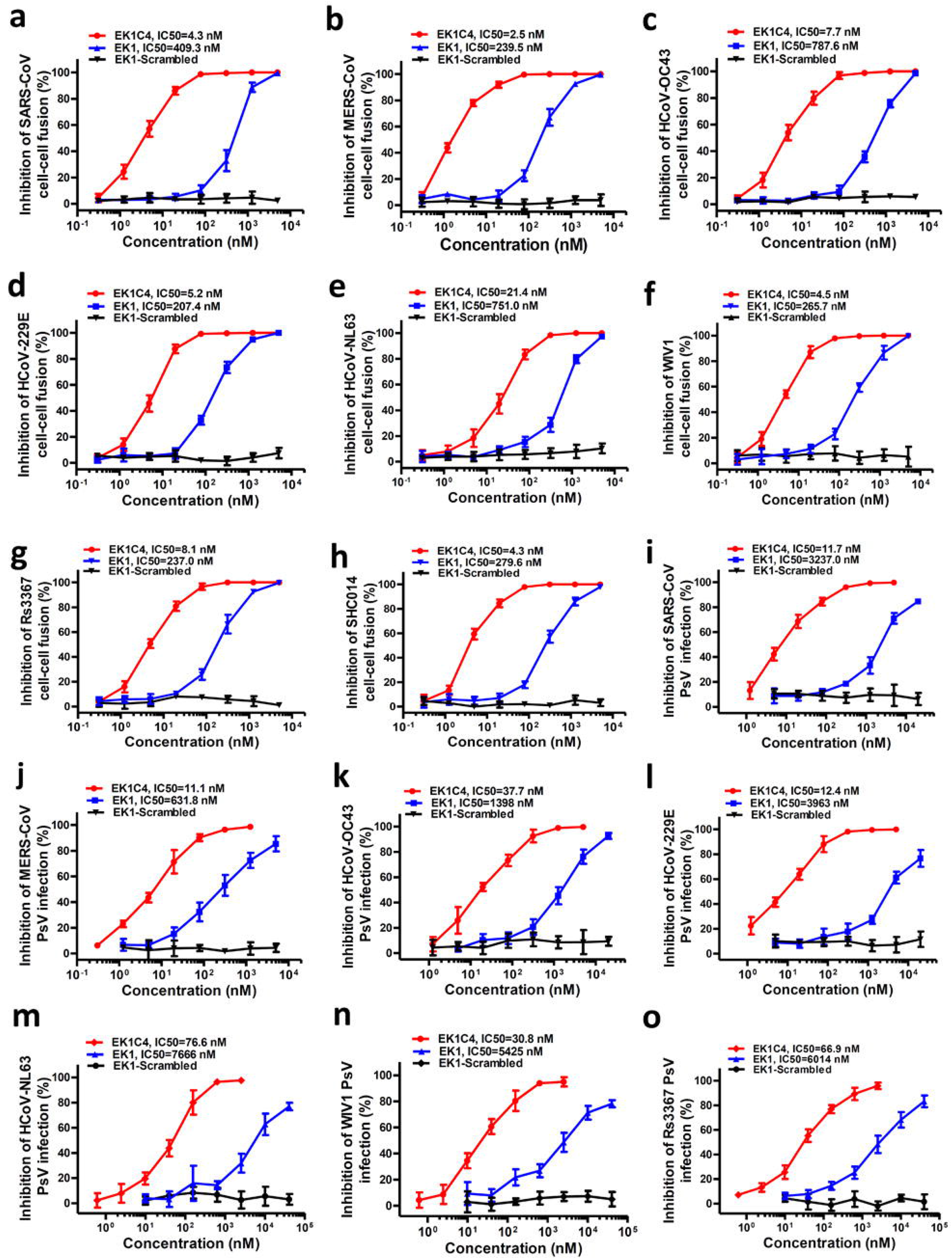
EK1C4 broadly and potently inhibited cell-cell fusion and PsV infection mediated by S protein of divergent HCoVs. **a** to **h**. Inhibitory activity of EK1C4 in cell-cell fusion mediated by the S proteins of SARS-CoV (**a**), MERS-CoV (**b**), HCoV-OC43 (**c**), HCoV-229E (**d**), HCoV-NL63 (**e**), WIV1 (**f**), Rs3367 (**g**) and SHC014 (**h**). **i** to **o**. Inhibitory activity of EK1C4 in PsV infection assays against SARS-CoV (**i**), MERS-CoV (**j**), HCoV-OC43(**k**), HCoV-229E (**l**), NL63 (**m**), WIV1 (**n**) and Rs3367 (**o**). Experiments were repeated twice, and the data are expressed as means ± SD.

We also assessed the antiviral activity of EK1C4 on PsV infection by divergent coronaviruses. As expected, EK1C4 showed much more potent activity than EK1 (IC_50_ ranging from 631.8 to 3237.0 nM) against SARS-CoV, MERS-CoV, and HCoV-OC43 infection with IC_50_ of 11.7 nM, 11.1 nM and 37.7 nM, respectively (Fig. 5i-k). EK1C4 also effectively blocked PsV infection of α-HCoVs, including HCoV-229E and HCoV-NL63, with IC_50_ of 12.4 nM and 76.6 nM, respectively, which was about 319- and 99-fold more active than EK1 (IC_50_ ranging from 3,963 to 7,666 nM) (Fig. 5l-m). Similarly, by cholesteryl modification with “GSGSG-PEG4” linker, the inhibitory activity of EK1 could be significantly increased on PsV infection from SL-CoVs, including WIV1 and Rs3367, where EK1C4 showed potent inhibitory activity with IC_50_ of 30.8 nM and 66.9 nM, respectively, which is 175-fold to 89-fold more potent than that of EK1 (Fig. 5n-o).

### EK1C4 possesses the most potent inhibitory activity against *in vitro* infection by live coronaviruses

We further assessed the inhibitory activity of EK1C4 against live HCoVs infection, including SARS-CoV-2, MERS-CoV, HCoV-OC43, HCoV-229E, and HCoV-NL63. Importantly, EK1C4 effectively blocked SARS-CoV-2 infection at the cellular level in a dose-dependent manner with IC_50_ of 36.5 nM, being 67-fold more active than that of EK1 (IC_50_=2,468 nM) (Fig. 6a), which is consistent to the results of cell-cell fusion assay and PsV infection assay mediated by SARS-CoV-2 S protein. Similarly, EK1C4 also showed more potent antiviral activity than EK1 against MERS-CoV, HCoV-OC43, HCoV-229E, and HCoV-NL63 infection with IC_50_s of 4.2 nM, 24.8 nM, 101.5 nM and 187.6 nM, respectively, which are 190-, 62-, 42- and 19-fold more potent than those of EK1, respectively (Fig. 6b-e). We next assessed the cytotoxicity of EK1C4 on various target cells and found that the half cytotoxic concentration (CC_50_) was beyond 5 μM, which is the highest detection concentration of EK1C4 (Fig. S3). Therefore, the selectivity index (SI=CC_50_/IC_50_) of EK1C4 is >136, suggesting that EK1C4 is a promising SARS-CoV-2 fusion inhibitor with little, or even no, toxic effect *in vitro*. Further, we explored the potent antiviral mechanism of EK1C4 and found that the complexes of EK1C4/SARS-2HR1, EK1C4/MERS-HR1, and EK1C4/SARS-2HR1 harbor higher stability and increased *Tm* values than those of the complexes formed by EK1 and HR1s (Fig. S4). These results suggested that increased antiviral activity of EK1C4 should be related its increased binding affinity with HR1, but their detailed interactions require further studies.

**Fig. 6.**
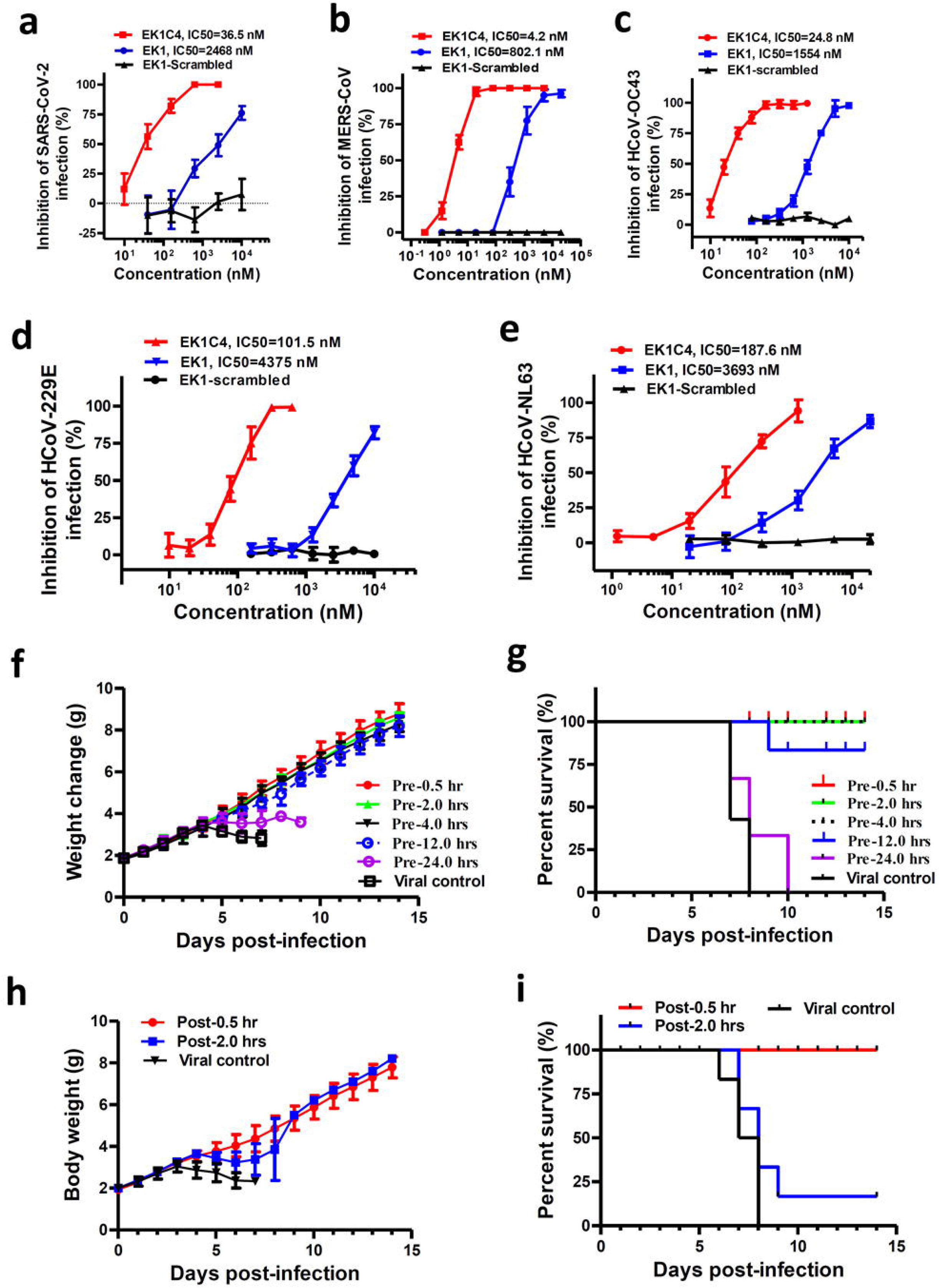
EK1C4 effectively inhibited live-CoVs infection *in vitro* and *in vivo.* **a-e.** Inhibitory activity of EK1 on live HCoV replication for SARS-CoV-2 (**a**), MERS-CoV (**b**), HCoV-OC43 (**c**), HCoV-229E (**d**), and HCoV-NL63 (**e**). **f-g.** *In vivo* prophylactic efficacy of EK1C4 against HCoV-OC43 infection in mice. Body weight change (**f**) and survival curves (**g**) of mice challenged with HCoV-OC43. **h-i.** *In vivo* therapeutic efficacy of EK1C4 against HCoV-OC43 infection in mice. Body weight change (**h**) and survival curves (**i**) of mice challenged with HCoV-OC43. Experiments were repeated twice, and the data are expressed as means ± SD.

### Intranasally applied EK1C4 showed strong protection of mice against HCoV-OC43 infection

Recently, SARS-CoV-2 rapidly spread in humans by transmitting through the respiratory tract. Here, we used an HCoV-OC43 infection mouse model to further investigate the potential prophylactic effect of EK1C4 in clinical applications *via* the intranasal administration route (Fig. 6f-g). In the OC43-infected mouse model, we treated newborn mice with EK1C4 at a single dose of 0.5 mg/kg 0.5 h (Pre-0.5), 2 h (Pre-2), 4 h (Pre-4), 12 (Pre-12) and 24 h (Pre-24) before challenging with HCoV-OC43 at 100 TCID_50_ (50% tissue culture infectious dose). Starting from 4 days’ post-infection (dpi), the body weight of mice in the viral control group decreased significantly along with 100% mortality (Fig. 6f-g). The final survival rates of mice in Pre-0.5, Pre-2, Pre-4, Pre-12 and Pre-24 groups were 100%, 100%, 100%, 83% and 0%, respectively (Fig. 6f-g). In contrast, EK1 with a single dose of 20 mg/kg via nasal administration exhibited very promising prophylactic effect in the Pre-0.5 h and Pre-1 h groups, whereas all mice in the EK1-Pre-2 h group eventually died similarly to the mice in the viral control group (Fig. S5). These results suggested that EK1C4 has better stability, antiviral activity, and prolonged half-life in the airway environment when compared with EK1.

We then tested the therapeutic effect of EK1C4 0.5 h (Post-0.5 group) and 2 h (Post-2 group) after HCoV-OC43 infection (Fig. 6h-i). The Post-0.5 group and Post-2 group mice showed 100% and 16.7% survival rate, respectively, suggesting that EK1C4 harbors good therapeutic effect after a short period of HCoV-OC43 infection, possibly resulting from the establishment of HCoV-OC43 infection in mouse brain where EK1C4 cannot get through the blood brain barrier via nasal administration^14^. As shown in Fig. S6, high viral titer was detected in brains of all 5 mice in Pre-24 group and 4 out of 5 mice in Post-2 group, but was not detected in brain tissues of all mice in Pre-0.5, Pre-2, Pre-4, and Post-0.5 groups, while only moderate level of viral titer was detected in brain tissue in one of the 5 mice in Pre-12 group (Fig. S6 a and b). Similar to those in the viral control mice, mice in Pre-24 and Post-2 groups exhibited similar histopathological changes in brain tissues, including vacuolation, degeneration, and infiltration. However, the brain tissues of mice in Pre-0.5, Pre-2, Pre-4, Pre-12 and Post-0.5 group as well as the normal control group showed no apparent histopathological changes (Fig. S6 c).

## Discussion

Over the past 20 years, highly infectious pathogens have been emerging increasingly, such as SARS-CoV in 2003 and MERS-CoV in 2012 ^20–22^. In 2018, WHO proposed “Disease X” in the blueprint priority diseases for any new unknown pathogen that may cause an epidemic or pandemic in the future, calling for the development of effective and safe vaccines and antivirals to prevent and treat such Disease X. Indeed, at the end of 2019, the outbreak of Wuhan pneumonia with an unknown etiological agent, the first Disease X following WHO’s announcement was reported to WHO. Shortly thereafter, a novel coronavirus, SARS-CoV-2 (also known as 2019-nCoV or HCoV-19), was identified to be the etiology of the Wuhan pneumonia, *i.e.*, COVID-19 as designated by WHO.

Unlike SARS-CoV, live SARS-CoV-2-infected cells were found to form typical syncytium, suggesting that SARS-CoV-2 may mainly utilize the plasma membrane fusion pathway to enter and replicate inside host cells. Consistently, in the cell-cell fusion system, SARS-CoV-2 S protein could effectively mediate the formation of syncytium between the effector cell and the target cell in the absence of an exogenous proteolytic enzyme, e.g., trypsin, while SARS-CoV S protein could not. Actually, the plasma membrane fusion pathway is more efficient than the endosomal membrane fusion pathway for most viruses because the latter is more prone to activating the host cell antiviral immunity^23,24^. Generally, β-B coronaviruses lack the S1/S2 furin-recognition site, and their S proteins are uncleaved in the native state. For example, SARS-CoV enters into the cell mainly *via* the endosomal membrane fusion pathway where its S protein is cleaved by endosomal cathepsin L and activated^25^. Inducing the S1/S2 furin-recognition site could significantly increase the capacity of SARS-CoV S protein to mediate cellular membrane surface infection ^26^. Interestingly, SARS-CoV-2 harbors the S1/S2 cleavage site in its S protein, but its specific role in S protein-mediated membrane fusion and viral life-cycle remains to be further explored (Fig. S7). A recent report suggested that SARS-CoV-2 mainly used TMPRSS2 for plasma membrane fusion; this means that the TMPRSS2 inhibitor might constitute an option for blocking SARS-CoV-2 fusion with and entry into the host cell ^27^.

The 6-HB structure formed by HR1 and HR2 regions in the S2 subunit of HCoVs plays a key role during the viral membrane fusion process, which makes it one of the most important targets for drug design. In previous studies, we have found that HR1 and HR2 of SARS-CoV-2 could form a stable coiled-coil complex, but the detailed conformations remain unknown. According to the X-ray crystallographic analysis of the complex formed by HR1 and HR2 of SARS-CoV-2 (Fig. 2b), it is a typical 6-HB fusion core structure similar to those of SARS-CoV and MERS-CoV. Although the amino acid sequences of HR2 domain from SARS-CoV and SARS-CoV-2 are fully identical, multiple residue differences occur in the HR1 domain of SARS-CoV-2. However, instead of weakening the interaction between HR1 and HR2, such unilateral difference seems to form new interactions in some regions and enhance the existing ones in other regions (Fig. 3). When K991 in SARS-CoV HR1 was replaced with S929 in SARS-CoV2 HR1, a new, strong hydrogen bond was formed with a distance of 2.4 Å. K933 forms a new interaction with N1192 in SARS-CoV-2 with a distance of 2.7 Å, whereas the corresponding position in SARS-CoV has no such interaction. In the other two regions, E918 binds to R1166 and K929 binds to E1163 in SARS-CoV, both of which were enhanced in SARS-CoV-2. These results suggest that this new HCoV has evolved with improved binding affinity between HR1 and HR2 domains, which may accelerate the viral membrane fusion process and enhance viral infectivity or transmissibility. A recent study also found that the binding affinity between ACE2 receptor on the host cell and RBD in S protein of SARS-CoV-2 is more than 10-fold higher than that of SARS-CoV, which may also be associated with the increased infectivity and transmissibility of SARS-CoV-2^12^.

The conjugation of cholesterol to viral entry inhibitor has been proved to be an effective strategy to enhance the antiviral activity, such as C34 peptide for HIV-1 ^28^. However, the mechanism of this enhancement, especially the role of cholesterol group in the C-terminal tail of entry inhibitor, is still unclear. There is a possibility that the cholesterol group could anchor to the target membrane to facilitate the binding of inhibitor to the HR1 targets. However, we noticed that binding affinity between EK1C4 and SARS-CoV-2-HR1P is significantly enhanced than EK1 peptide alone, which suggested that cholesterol group may be involved in binding to HR1P directly (Fig. S4). Therefore, using structural simulation and docking method, we predicted a possible model of EK1C4 in binding with SARS-CoV-2 HR1P (Fig. S8). In this model, the EK1C4 peptide anchors to one of the three hydrophobic grooves of HR1 trimer via its EK1 moiety, and also anchors to another adjacent hydrophobic groove of HR1 trimer via its cholesterol moiety. The cholesterol group of EK1C4 may bind to HR1P through hydrophobic interactions, while several hydrogen bonds may form between HR1 and helical region of EK1C4. The intermediated GSGSG-PEG4 linker of EK1C4 peptide is just enough to connect these two moieties on the two binding targets. Admittedly, the exact mechanism and structure of EK1C4 need more studies in the future.

In the past few decades, the viral HR1 domain has been proved to be an important target for the development of viral fusion and entry inhibitors. In the early outbreak of MERS, we quickly solved the 6-HB fusion core structure formed by MERS-CoV S protein HR1 and HR2 domains and designed the fusion inhibitory peptide HR2P-M2 which proved to be highly effective in blocking its spike protein-mediated membrane fusion and inhibit *in vitro* MERS-CoV infection^16^. The results from animal experiments showed that intranasal application of HR2P-M2 peptide could effectively protect mice from MERS-CoV infection with reduction of virus titers in the lung more than 1000-fold^29^. However, the MERS-CoV HR2P-M2 peptide could not inhibit SARS-CoV infection, suggesting that this peptide lacks cross-inhibitory activity against other β-CoVs, such as SARS-CoV and bat SARSr-CoVs. To be well prepared for combating the emerging coronaviruses with epidemic or pandemic potential, we designed and synthesized the first pan-coronavirus fusion inhibitor, EK1, and found that EK1 exhibited potent inhibitory activity against all HCoVs that we tested, including SARS-CoV and MARS-CoV, as well as bat SARSr-CoVs. As expected, we recently have shown that EK1 is also effective in inhibiting infection of the novel β-CoV, SARS-CoV-2 ^15^. We then optimized EK1 peptide in hopes of improving its fusion inhibitory activity. Indeed, we found that one of the modified EK1 peptides, EK1C4, was 226-fold and 149-fold more potent against SARS-CoV-2 S protein-mediated membrane fusion and PsV infection, respectively, than EK1. EK1C4 also showed broad-spectrum inhibitory activity against infection by SARS-CoV, MERS-CoV and other HCoVs. EK1C4 showed prolonged and significant prophylactic effect against HCoV-OC43 infection in mouse model, suggesting that EK1C4 may also be used as an inhibitor against SARS-CoV-2 infection *in vivo*. Consistent with other studies ^35^, HCoV-OC43 was showed as a typical neurotropic virus in the mouse model, and quickly entered and established infection in mouse brain tissue, leading to the relatively weak therapeutic effect of EK1C4 via intranasal administration. However, SARS-CoV-2 mainly infected and caused severe pathological changes in human lung tissue ^4^. Therefore, EK1C4 administered intranasally is expected to have good therapeutic potential against SARS-CoV-2 infection.

Currently, no specific anti-CoV therapeutics or prophylactics have been used in clinics for treatment or prevention of SARS-CoV-2 infection. A number of nonspecific antiviral drugs, including IFN, lopinavir-ritonavir (HIV protease inhibitors), chloroquine, favipiravir (T-705) and remdesivir (GS-5734), have been used in clinics in China to treat SARS-CoV-2 infection ^30^. Their *in vivo* efficacies still require further confirmation. Their potential use for treatment of infection by other coronaviruses and emerging coronaviruses in the future is unclear. Compared with these clinically used nonspecific antiviral drugs, EK1C4 has more advantages for treatment and prevention of SARS-CoV-2 infection. First, the sequence of its target, the HR1 domain in S2 subunit of S protein, is highly conserved. Therefore, EK1C4 possesses a high genetic barrier to resistance and cannot easily induce drug-resistant mutations. Second, EK1C4 can be used in an intranasal formulation to prevent coronavirus infection. The small bottles can be carried easily by persons who will have close contact with infected patients or high-risk populations. Third, EK1C4 can be used in inhalation formulation for treatment of patients to reduce the viral loads in their lungs, thus attenuating the acute lung injury caused by viral infection and reducing the chance to spread the virions to the closely contacted persons. The inhalation equipment can be used in home or hotel room, reducing the expense of staying in hospitals. Fourth, EK1C4 is expected to be safe to humans because it will be used locally, not systemically, and peptide drugs are generally safer than chemical drugs. Fifth, because of its broad-spectrum anti-coronavirus activity, EK1C4 can be used for treatment and prevention of infection by not only SARS-CoV-2, but also other HCoVs. Sixth, recently 103 SARS-CoV-2 genomes have been identified ^31^, but we found that both the HR1 and HR2 domains among those reported genomes show 100% identity (Fig. S9), indicating the high conservation of EK1C4 target. In the meantime, the HR2 derived peptides have much larger interface on HR1 domain, making it more resistant to the viral mutations. Therefore, EK1C4 shows exceptional promise to be developed as the first pan-CoV fusion inhibitor-based antiviral therapeutic or prophylactic for treatment or prevention of infection by the currently circulating SARS-CoV-2 and MERS-CoV and the future reemerging SARS-CoV and emerging SARSr-CoVs.

## Methods

### Cell Lines, viruses and Peptides

The human primary embryonic kidney cell line (293T) (CRL-3216™), Calu-3 (HTB-55™), A549 (CCL-185), Vero E6 (CRL-1586™), RD (CCL-136™), and LLC-MK2 Original (CCL-7™) cells were obtained from the American Type Culture Collection (ATCC). Human hepatoma Huh-7 cells were from the Cell Bank of the Chinese Academy of Sciences (Shanghai, China), and 293T cells stably expressing human ACE2 (293T/ACE2) cells were kindly provided by Dr. Lanying Du. All of these cell lines were maintained and grown in Dulbecco’s Modified Eagle’s Medium (DMEM, Invitrogen, Carlsbad, CA, USA) containing 100 U/ml penicillin, 100 mg/ml streptomycin, and 10% heat-inactivated fetal calf serum (FCS) (Gibco).

Patient-derived COVID-19 (BetaCoV/Wuhan/WIV04/2019) was isolated by the Wuhan Institute of Virology ^6^. MERS-CoV-EMC/2012 was originally provided by Chuan Qin (Beijing Key Laboratory for Animal Models of Emerging and Re-emerging Infectious Diseases). ATCC strain of Human coronavirus 229E (HCoV-OC43, VR-740), as well as Human coronavirus OC43 (HCoV-229E, VR-1558) and HCoV-NL63 (Amsterdam strain) strains were amplified in Huh-7, HCT-8 and LLC-MK2 cells, respectively.

Peptides were synthesized by Chao Wang (Beijing Institute of Pharmacology and Toxicology). The sequences of EK1 (SLDQINVTFLDLEYEMKKLEEAIKKLEESYIDLKEL) and EK1-scrambled (LKVLLYEEFKLLESLIMEILEYQKDSDIKENAEDTK) have been reported in our previous study ^14^

### Plasmids

The envelope-expressing plasmids of SARS-2-S (pcDNA3.1-SARS-2-S), SARS-S (pcDNA3.1-SARS-S), MERS-S (pcDNA3.1-MERS-S), OC43-S (pcDNA3.1-OC43-S), NL63-S (pcDNA3.1-NL63-S), 229E-S (pcDNA3.1-229E-S), and bat SARS-like CoV-S (pcDNA3.1-WIV1-S, pcDNA3.1-Rs3367-S and pcDNA3.1-SHC014-S), and the plasmids pAAV-IRES-EGFP that encode EGFP as well as the luciferase reporter vector (pNL4-3.Luc.R-E-) were maintained in our laboratory.

### Cell–cell fusion assay

The establishment and detection of several cell–cell fusion assays are as previously described ^14,16^. In brief, Huh-7 cells (for testing all coronaviruses) or 293T/ACE2 cells (for testing SARS-CoV-2) were used as target cells. For preparing effector cells expressing S protein a coronavirus, 293T cells were transfected with one of the S protein expression vectors, including 293T/SARS-CoV-2/GFP, 293T/MERS-CoV/GFP, 293T/HCoV-229E/GFP, 293T/SARS-CoV/GFP, or 293T/SL-CoV/GFP, 293T/HCoV-OC43/GFP, 293T/HCoV-NL63/GFP or empty plasmid pAAV-IRES-EGFP. For SARS-CoV S-, SL-CoV S-, OC43 S- or NL63 S-mediated cell-cell fusion assays, effector cells and target cells were cocultured in DMEM containing trypsin (80 ng/mL) for 4 h, while for SARS-CoV-2 and MERS-CoV S-mediated cell-cell fusion assays, effector cells and target cells were cocultured in DMEM without trypsin but 10% FBS for 2 h. After incubation, five fields were randomly selected in each well to count the number of fused and unfused cells under an inverted fluorescence microscope (Nikon Eclipse Ti-S).

### Inhibition of HCoV S-mediated cell-cell fusion

The inhibitory activity of a peptide on a HCoV S-mediated cell-cell fusion was assessed as previously described^14,16^. Briefly, a total of 2×10^4^ cells/well target cells (Huh-7) were incubated for 5 h. Afterwards, 10^4^ cells/well effector cells (293T/S/GFP) were added in the presence or absence of a peptide at the indicated concentrations at 37 °C for 2 h. 293T/EGFP cells with phosphate-buffered saline (PBS) were used as a negative control. The fusion rate was calculated by observing the fused and unfused cells using fluorescence microscopy.

### Inhibition of pseudotyped HCoV infection

293T cells were cotransfected with pNL4-3.luc.RE (the luciferase reporter-expressing HIV-1 backbone) and pcDNA3.1-SARS-CoV-2-S (encoding for CoVs S protein) using VigoFect (Vigorous Biotechnology, Beijing, China) ^16,32,33^. Pseudotyped particles were efficiently released in the supernatant. The supernatant was harvested at 72 h post-transfection, centrifuged at 3000× g for 10 min, and frozen to −80 °C. To detect the inhibitory activity of a peptide on infection of coronavirus PsV, target cells (293T/ACE2 for SARS-CoV-2, SARS-CoV and SL-CoVs; RD cells for HCoV-OC43; Huh-7 for other CoVs) were plated at a density of 10^4^ cells per well in a 96-well plate one day prior to infection ^14^. PsV was mixed with an equal volume of a peptide which was series diluted with PBS at 37 °C for 30 min. The mixture was transferred to the Huh-7 cells. Medium was changed after 12 h and incubation continued for 48 h. Luciferase activity was analyzed by the Luciferase Assay System (Promega, Madison, WI, USA).

### Inhibition of live HCoV replication

The inhibition assay for live SARS-CoV-2 and MERS-CoV was performed in a biosafety level 3 (BSL3) facility at the Wuhan Research Institute and Beijing Key Laboratory for Animal Models of Emerging and Re-emerging Infectious Diseases, respectively^6^. Inhibition activity of peptides on SARS-CoV-2 and MERS-CoV was determined by plaque reduction assay. Peptides with different dilution concentrations were mixed with SARS-CoV-2 (100 TCID50) for 30 minutes and then added to monolayer VERO-E6 cells. After adsorption at 37 °C, the supernatant was removed, and 0.9 % methyl cellulose was overlaid on the cells. After 72 h, the plates were fixed and stained. Plaques were counted by fixing with 4% paraformaldehyde and staining with 0.1% crystal violet. To test the effect of peptide on HCoV-OC43, HCoV-229E and HCoV-NL63 replication, 50 μL of 100 TCID50 virus were mixed with an equal volume of peptide and incubated at 37 ° C for 1 hour. Afterwards, the mixture was added to RD, Huh-7 and LLC-MK2 cells, respectively. Cell Counting Kit-8 (CCK8, Dojindo, Kumamoto, Kyushu, Japan) assay was applied to determine cytopathic effect.

### Circular dichroism spectroscopy

The peptides or peptide mixtures were dissolved in PBS to prepare a solution with a final concentration of 10 μM at 37 °C for 30 min and then measured on a Jasco-815-circular dichroism spectrometer ^34^. The scanning wavelength range was 198-260 nm. Thermal denaturation detection starts at 222 nm with a 5 °C/min thermal gradient detection.

### Mouse infection studies

Newborn mice were bred from pregnant mice purchased from the Animal Center of Fudan University, and all the related experiments were carried out in strict accordance with institutional regulations (approval number 20190221-070, approval date 21 February 2019). Each group had 12 3-day-old mice. To test the protective effect of peptides on HCoV-infected mice, EK1C4 (0.5 mg/kg), EK1 (20 mg/kg) in 2 μl 28% Hydroxypropyl-β-Cyclodextrin (HBC), or phosphate-buffered saline (PBS) solution, were administered intranasally 0.5, 1, 2, 4, 12, and 24 h before challenge, or 0.5 and 2 h after challenge. Then mice were challenged intranasally with HCoV at a dose of 10^2^ TCID50. For the viral control group, the same volume of 28% HBC or PBS was administered intranasally. In each group, six mice were randomly selected for euthanasia on day 5 after infection, then five mice for collecting and assessing the viral titer in mouse brain, one mouse for brain histological examination. Body weight and survival of the remaining six mice in each group were monitored for 14 days ^35^

### Cytotoxicity assay

Cytotoxicity of the peptides to the cells (Vero-E6, Huh-7, LLC-MK2 and RD cells) was tested by using the Cell Counting Kit-8 (CCK-8). Briefly, each cell type was seeded into the wells of a 96-well microtiter plate (10,000 per well) and incubated at 37 °C for 12-15 h, replacing medium with DMED containing EK1C4 at graded concentrations to culture at 37 °C for 2 days; CCK-8 solution (10 μL per well) was added, followed by an additional incubation for 4 h. The absorbance was measured at 450 nm.

### Expression and purification of fusion protein HR1-L6-HR2 of SARS-CoV-2

The coding sequences of HR1 (residues 910 to 988) and HR2 (residues 1162 to 1206) domains of SARS-CoV-2 S2 subunits were tandem linked though a 6-residue linker (L6: SGGRGG). The resulting sequences encoding the fused HR1-L6-HR2 protein were then cloned into a modified pET-28a vector containing a His_6_-SUMO tag upstream of the multiple cloning site. The recombinant construct was expressed in *Escherichia coli* BL21 (DE3). Cells were grown in lysogeny broth (LB) media supplemented with 50 μg/mL kanamycin at 37 °C and were induced with 1 mM IPTG for 12 h at 16 °C overnight. Cells were harvested by centrifugation at 4500 g for 10 min at 4 °C and were lysed by high-pressure homogenizer twice after resuspension in buffer containing 25 mM Tris–HCl, pH 8.0, and 200 mM NaCl. The fusion proteins were isolated by Ni-affinity chromatography, and the SUMO tag was removed by Ulp1 enzyme (1:100 w/w) cleavage. HR1-L6-HR2 protein was concentrated and gel-filtered on a 10/300 Superdex 75 (GE Healthcare) column. Peak fractions containing HR1-L6-HR2 trimer were pooled and concentrated to 20 mg/ml through centrifugation (EMD Millipore).

### Crystallization and structure determination

Crystals were obtained at 16 °C for 7 days using the hanging drop vapor diffusion method by mixing equal volume of protein solution (HR1-L6-HR2, 10 mg/mL) and reservoir solution (10% PEG8000, 200 mM zinc acetate, 0.1 M MES, pH 6.0). Then crystals were flash-frozen and transferred to liquid nitrogen for data collection. On the in-house (Institute of Biophysics, Chinese Academy of Sciences) X-ray source (MicroMax 007 generator (Rigaku, Japan)) combined with Varimax HR optics (Rigaku, Japan), HR1-L6-HR2 crystals at 100 K were diffracted to 2.9-Å resolution at a wavelength of 1.5418 Å. A native set of X-ray diffraction data was collected with the R-AXIS IV ++ detector (Rigaku, Japan) with an exposure time of 3 min per image and was indexed and processed using iMosflm ^36^. The space group of the collected dataset is P21. Molecular replacement was performed with PHENIX.phaser ^37^ to solve the phasing problem, using the SARS-CoV S protein core structure (PDB code 1WYY) as a search model. The final model was manually adjusted in COOT and refined with Refmac ^38^. Data collection statistics and refinement statistics are given in Table 1. Coordinates were deposited in the RCSB Protein Data Bank (PDB code: 6LXT). The interaction model of EK1C4 peptide and HR1 domains of SARS-nCoV-2 was predicted by SWISS-MODEL sever ^39^ using 6XLT as reference for EK1 moiety, and by Autodock 4 software ^40^ for cholesterol moiety (Fig. S8).

**Table 1.**
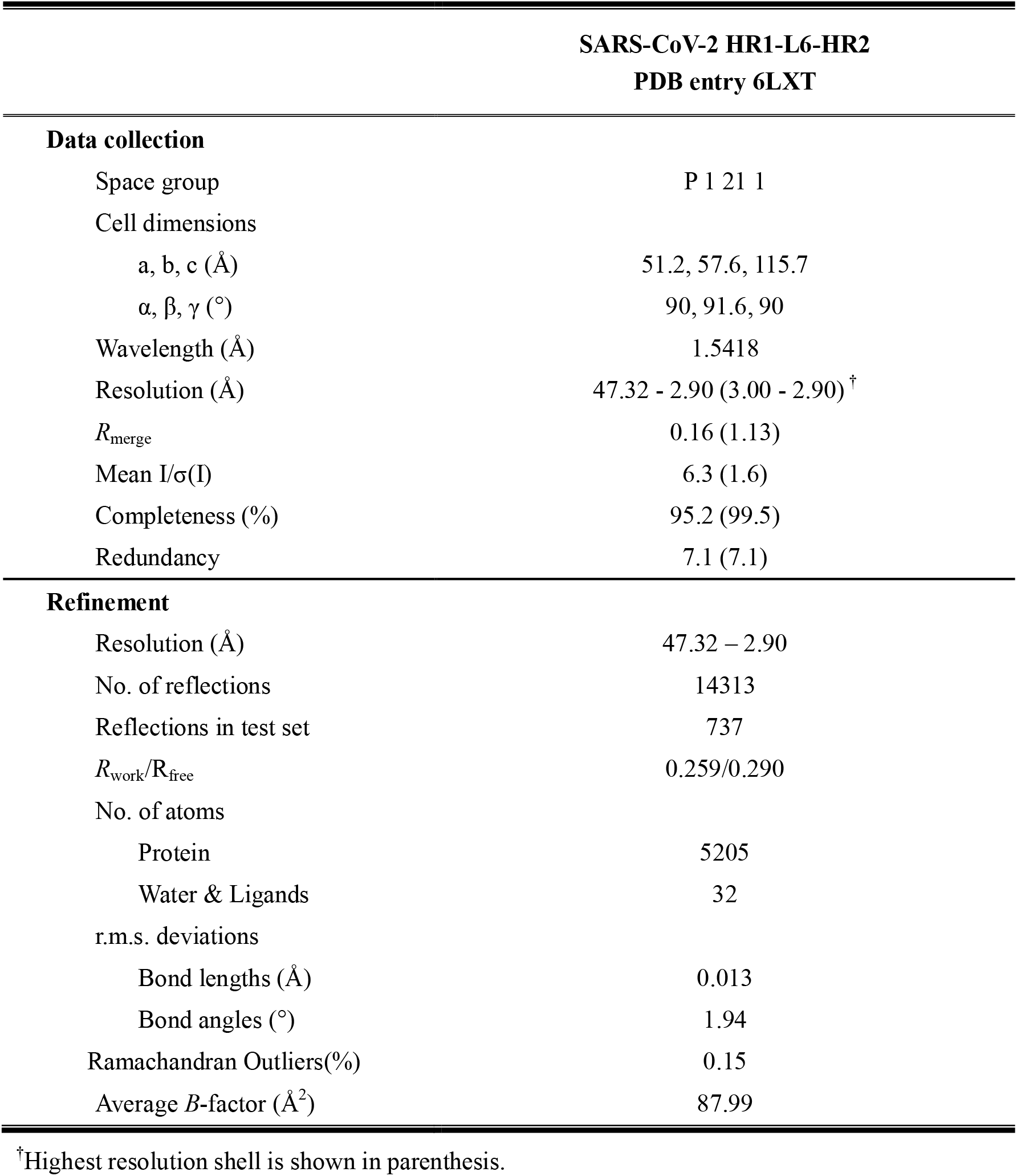
Data collection and refinement statistics.

### Statistical analysis

The survival rates of mice were analyzed by GraphPad Prism 5.0 software. CalcuSyn software was kindly provided by T.C. Chou, and the percent inhibition and IC_50_ values were calculated based on it ^34^.

