## Supplemental Figures for "Inhibition of SARS-CoV-2 infection (previously 2019-nCoV) by a highly potent pan-coronavirus fusion inhibitor targeting its spike protein that harbors a high capacity to mediate membrane fusion"

### Supplementary Fig. 1

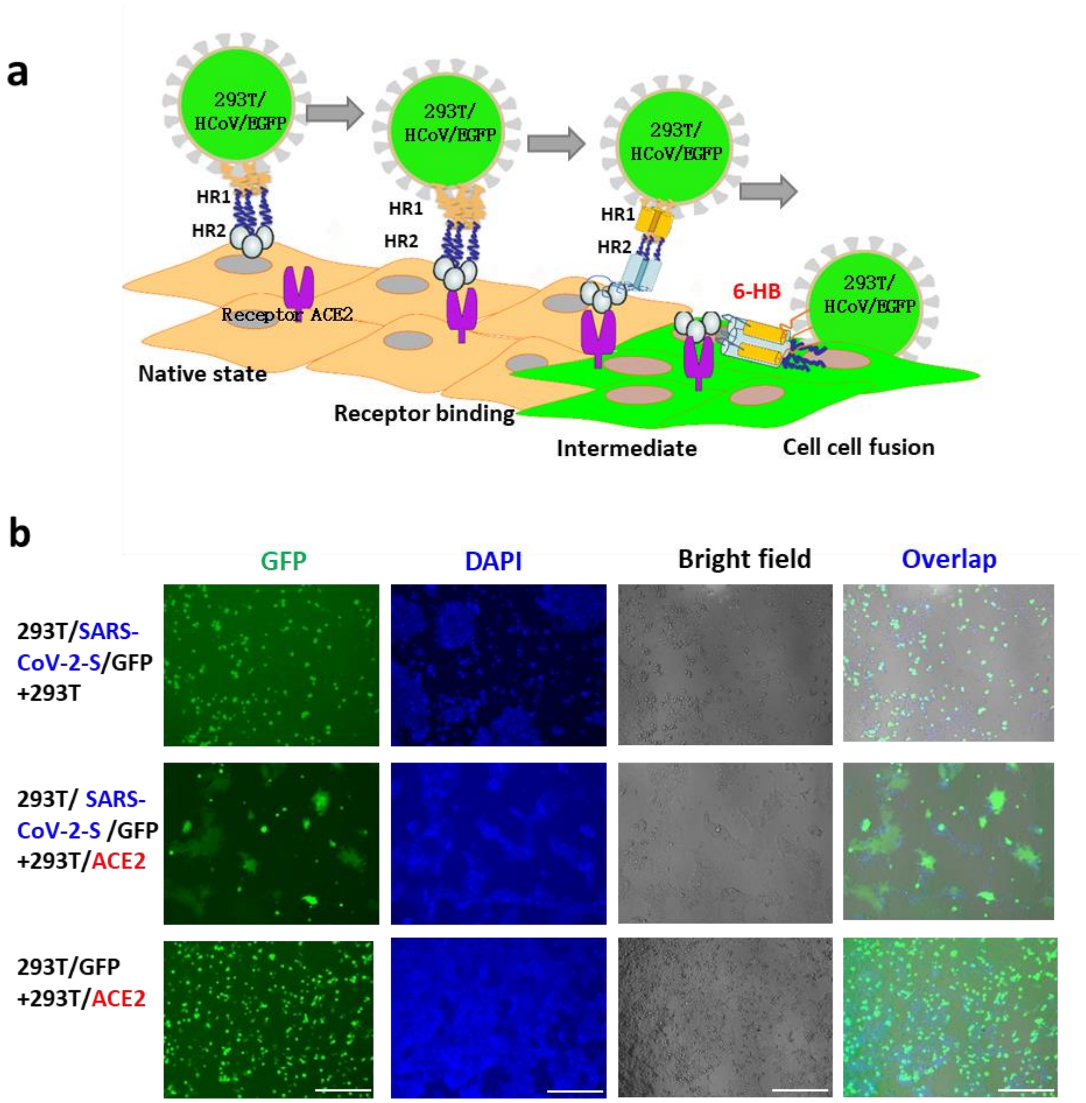


**Supplementary Fig. 1. Establishment of SARS-CoV-2 S-mediated cell-cell fusion.**

**a**. Schematic representation of SARS-CoV-2 S-mediated cell-cell fusion. **b**. Images of cell-cell fusion between 293T/SARS-CoV-2/EGFP cells and 293T cells (upper), 293T/SARS-CoV-2/EGFP cells and 293T/ACE2 cells (middle), 293T/EGFP cells and 293T/ACE2 cells (lower). Scale bar = 400 μm.

### Supplementary Fig. 2

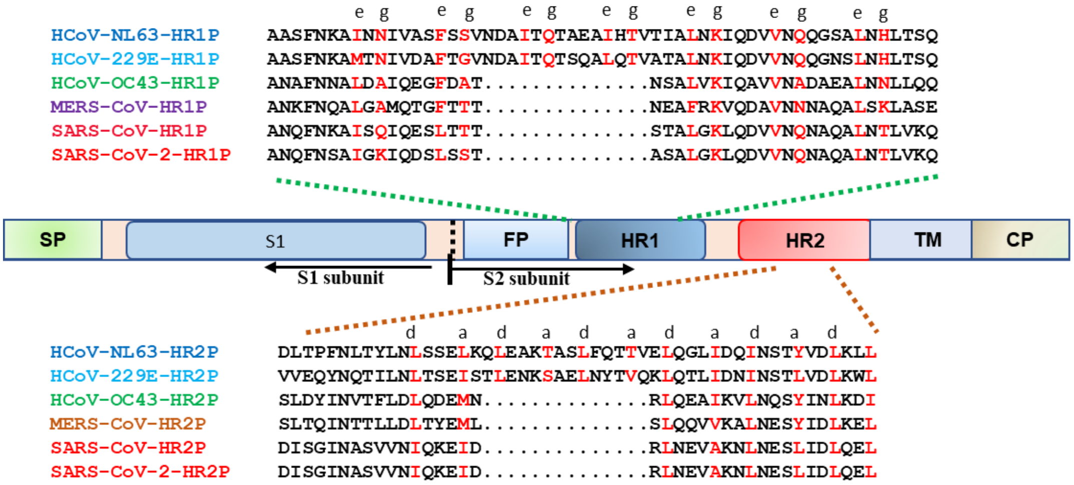


**Supplementary Fig. 2. Schematic representation of HCoV S protein and the sequences of the designed peptides (HR1Ps and HR2Ps).**

**Supplementary Fig. 3**

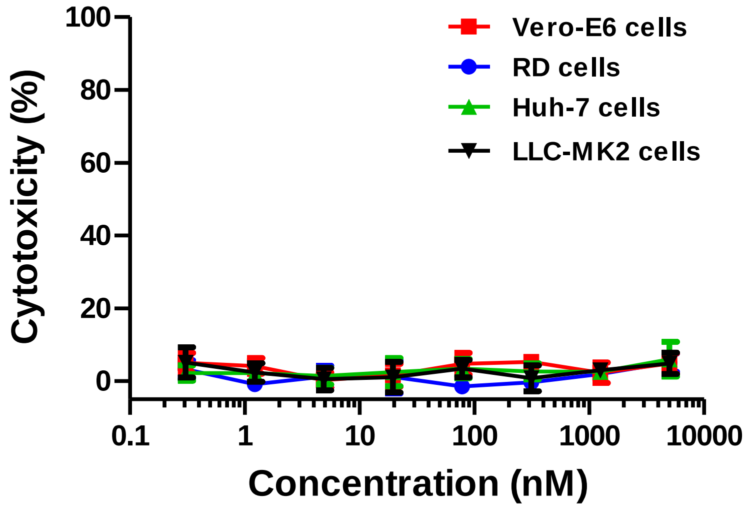


**Supplementary Fig. 3. Cytotoxicity of EK1C4 on Vero-E6, RD, Huh-7 and LLC-MK2 cells.**

### Supplementary Fig. 4

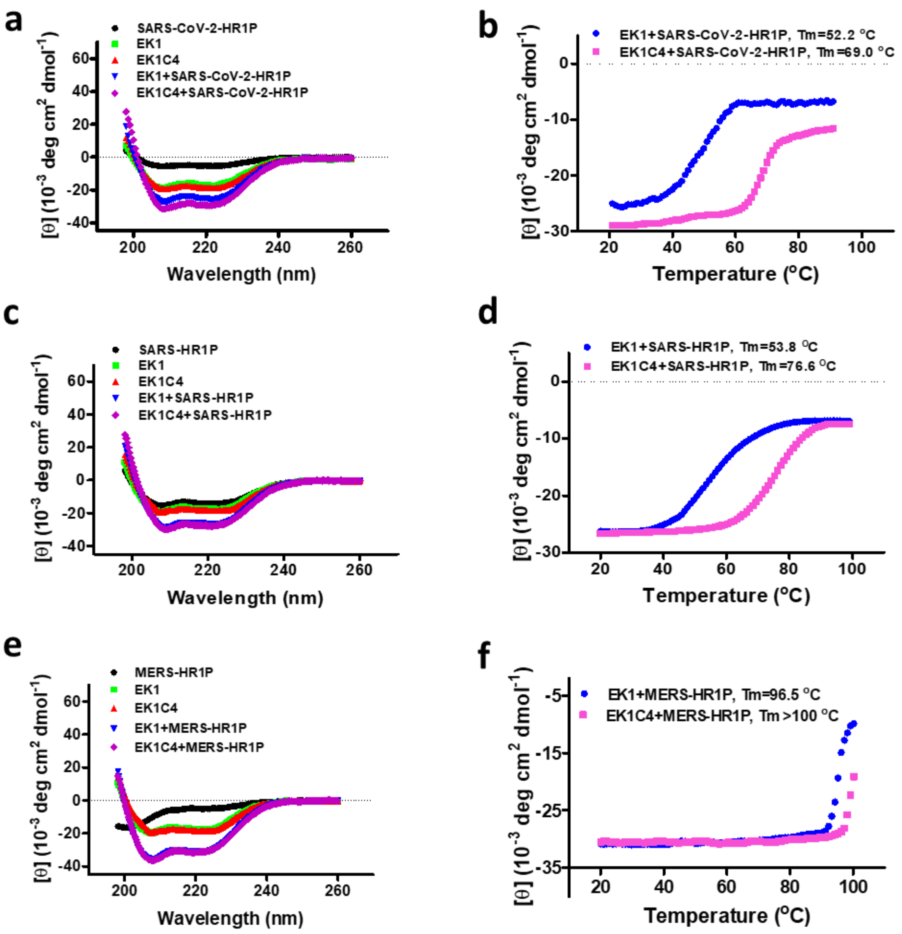


**Supplementary Fig. 4. Biophysical characterization of EK1C4 and its complexes with HR1Ps. a**. Circular-dichroism (CD) spectra for EK1, EK1C4, SARS-CoV-2-HR1P and their complexes in PBS. **b**. Melting curves of the complexes of EK1 / SARS-CoV-2-HR1P and EK1C4/SARS-CoV-2-HR1P. **c**. Circular-dichroism (CD) spectra for EK1, EK1C4, SARS-HR1P and their complexes in PBS. **d**. Melting curves of the complexes of EK1/SARS-HR1P and EK1C4/SARS-HR1P. **e**. Circular-dichroism (CD) spectra for EK1, EK1C4, MERS-HR1P and their complexes in PBS. **f**. Melting curves of the complexes of EK1/MERS-HR1P and EK1C4/MERS-HR1P.

### Supplementary Fig. 5

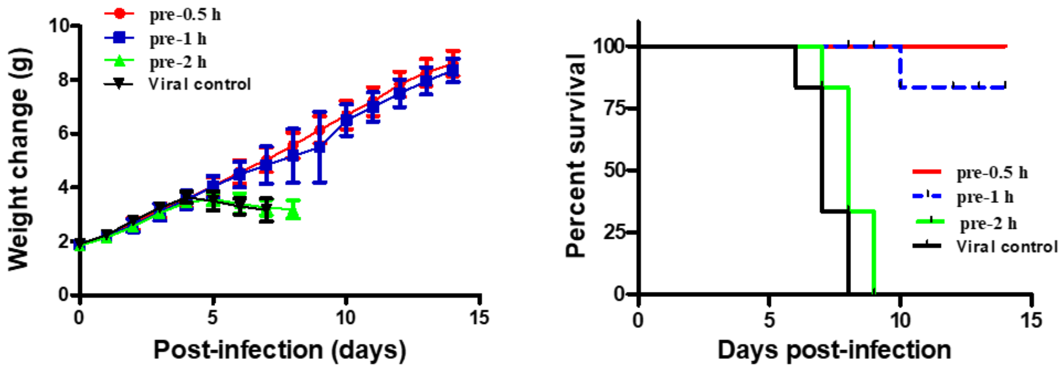


**Supplementary Fig. 5. *In vivo* prophylactic efficacy of EK1 against HCoV-OC43 infection in mice. a**. Body weight change of mice challenged with HCoV-OC43. **b**. Survival curves of mice challenged with HCoV-OC43.

### Supplementary Fig. 6

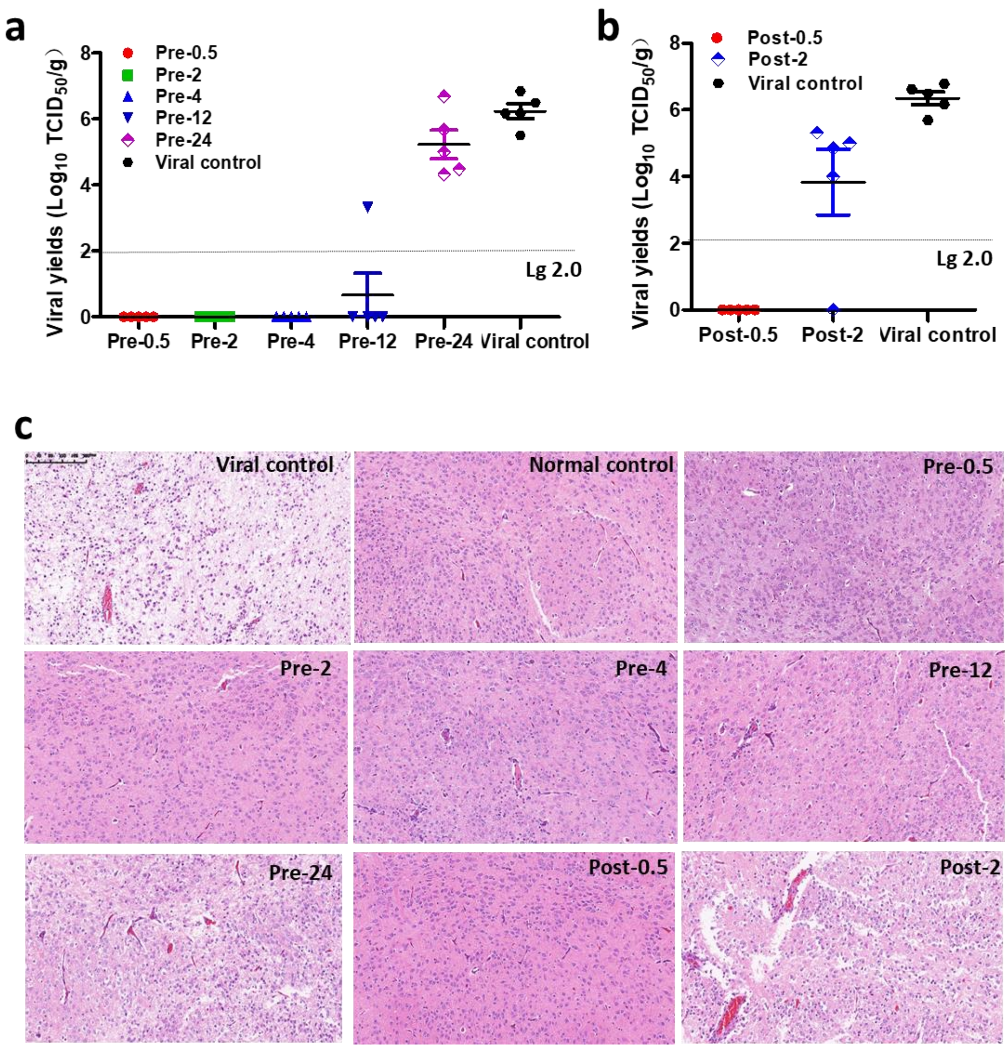


**Supplementary Fig. 6. The prophylactic and therapeutic effect of EK1C4 adminitered into mice before and after viral challenge, respectively.** a. Viral titer in brain tissues of mice in the viral control group and EK1C4-pre-treatment groups. b. Viral titer in brain tissues inthe viral control group and EK1C4-post-treatment groups. c. Histological examination of mouse brains..

### Supplementary Fig. 7

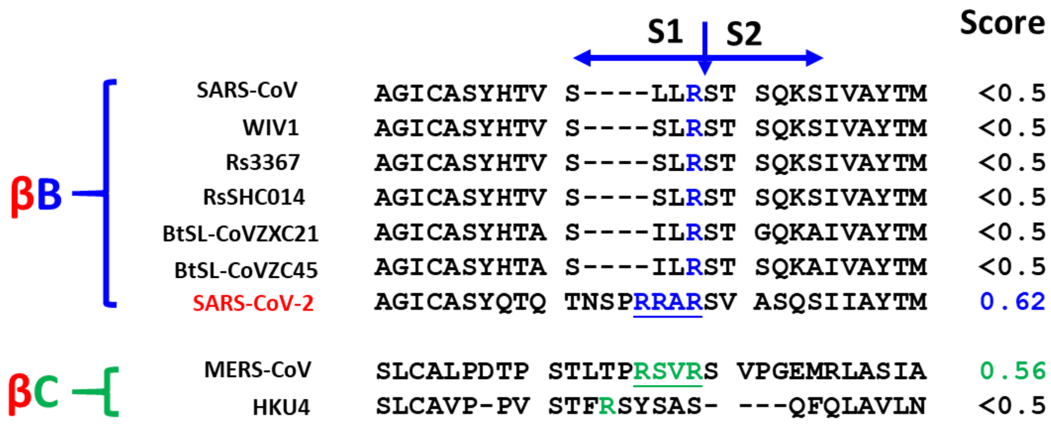


**Supplementary Fig. 7. Analysis of the S1/S2 furin-recognizable site in β-B and β-C coronaviruses.**

### Supplementary Fig. 8

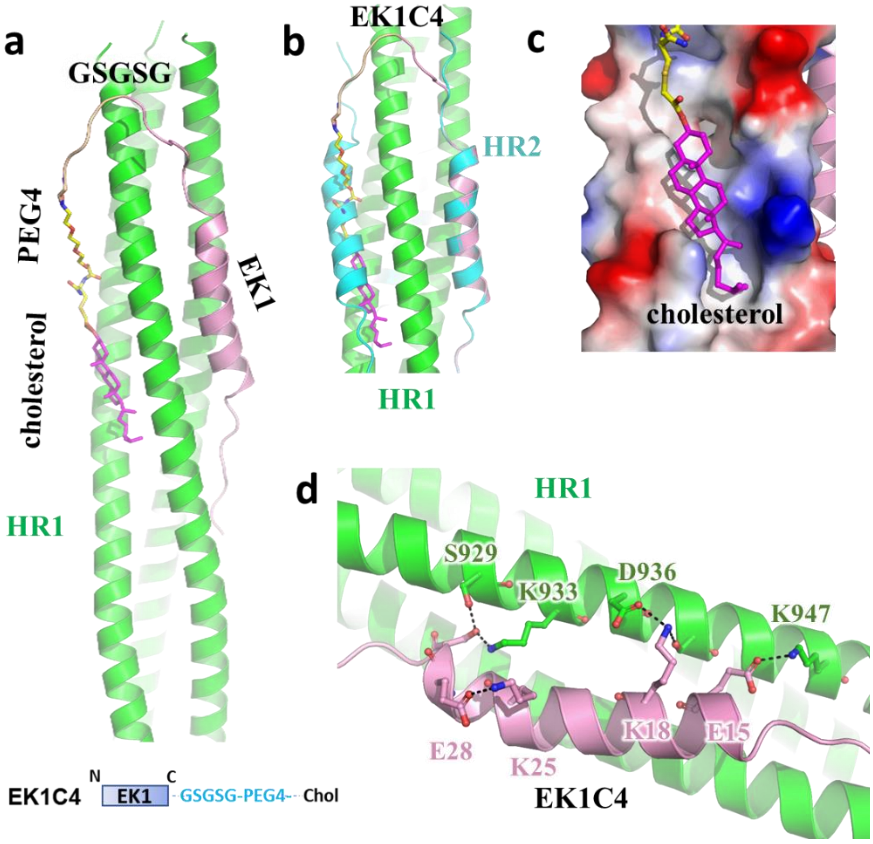


**Supplementary Fig. 8. The predicted model of interactions between EK1C4 peptide and HR1 domains of SARS-nCoV-2. a**. The interaction model of HR1 domain of SARS-nCoV-2 to EK1 peptide and cholesterol group are predicted by SWISS-MODEL sever using 6XLT as reference, and Autodock 4 software ， respectively. Structures are shown in cartoon representation, and each region of EK1C4 peptides is colored differently and labeled. **b**. The superposed structure of EK1C4 and HR2 peptides. **c**. The cholesterol domain is predicted to be buried in the hydrophobic grooves of SARS-CoV-2 HR1 domains. **d**. The EK1C4 peptide are predicted to interact with SARS-CoV-2 HR1 domain through several hydrogen bonds and salt bridges.

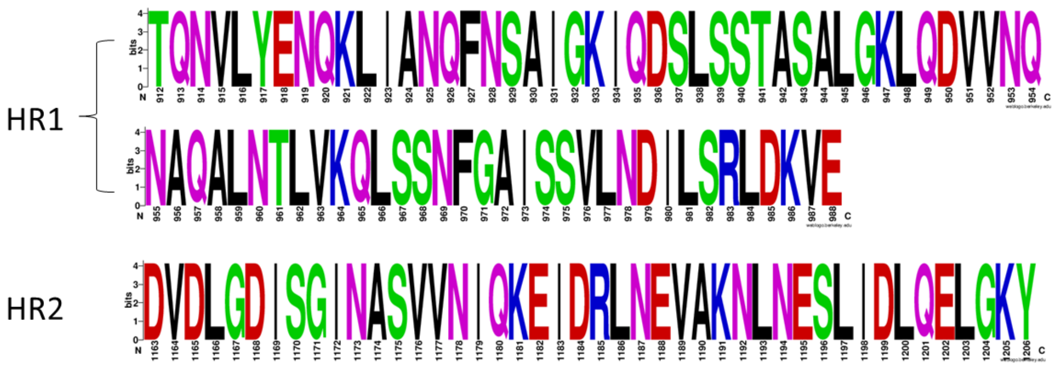


**Supplementary Fig. 9**

**Supplementary Fig. 9. The identical sequence of HR1 and HR2 domains in 103 SARS-CoV-2 genomes.**
